# A spatial atlas of inhibitory cell types in mouse hippocampus

**DOI:** 10.1101/431957

**Authors:** Xiaoyan Qian, Kenneth D. Harris, Thomas Hauling, Dimitris Nicoloutsopoulos, Ana B. Muñoz-Manchado, Nathan Skene, Jens Hjerling-Leffler, Mats Nilsson

## Abstract

Understanding the function of a tissue requires knowing the spatial organization of its constituent cell types. In the cerebral cortex, single-cell RNA sequencing (scRNA-seq) has revealed the genome-wide expression patterns that define its many, closely related cell types, but cannot reveal their spatial arrangement. Here we introduce *probabilistic cell typing by in situ sequencing* (pciSeq), an approach that leverages prior scRNA-seq classification to identify cell types using multiplexed *in situ* RNA detection. We applied this method to map the inhibitory neurons of hippocampal area CA1, a cell system critical for memory function, for which ground truth is available from extensive prior work identifying the laminar organization of subtly differing cell types. Our method confidently identified 16 interneuron classes, in a spatial arrangement closely matching ground truth. This method will allow identifying the spatial organization of fine cell types across the brain and other tissues.

---

The cortex contains many cell types, which differ in their spatial organization, morphology, connectivity, physiology, and gene expression. Although the diversity of cortical cells was known to classical neuroanatomists, the true complexity of cortical cells has only become clear since the development of transcriptomics^1–6^. Many cell types previously thought to be homogeneous in fact contain multiple fine subclasses, which share expression of the great majority of their genes, but are distinguished by relatively small sets of genes. Hippocampal area CA1 contains at least 20 subtypes of molecularly-distinct inhibitory interneurons, which are arranged in a stereotyped spatial organization^7–9^, and recordings have shown that even closely related cell types have different *in vivo* functions^10^. Analysis of CA1 inhibitory classes by scRNA-seq yields clusters strikingly consistent with these classically-defined types^5^. Mapping the spatial organization of CA1 interneurons is thus not only important to understand the brain’s memory circuits, but also provides a powerful way to validate spatial cell type mapping approaches, using the spatio-molecular ground truth provided by this system.

Here we provide a spatial map of CA1 interneuron types, using a new approach to *in situ* cell typing based on *in situ* RNA expression profiling. While several approaches to multiplexed *in situ* RNA detection and cell type classification have been proposed^11,12^ (e.g. *in situ* sequencing^13^, FISSEQ^14^, SNAIL^15^, MERFISH^16^, CorrFISH^17^, osmFISH^18^ and seqFISH^19^), none have yet shown the ability to distinguish fine cell types. The current method achieves this using three design criteria suggested by scRNA-seq analysis. First, for typing of closely related cells such as CA1 inhibitory neurons, of the order 100 genes suffices to classify cells accurately^5^. Second, false positive reads present much more of a problem for cell typing than low detection efficiency. Indeed, because RNA expression levels follow a negative binomial distribution^20^ (whose standard deviation scales with its mean), detecting just a few copies of a genuinely-expressed gene can confidently identify cell types, provided one can be sure these are not misdetections. Third, using the cell types revealed by previous large-scale scRNA-seq analysis to define types avoids needing to relearn cell classes from noisier *in situ* data. We show that this combination allows cell typing of closely-related neuronal classes, verified by the ground truth available from CA1’s laminar architecture.

Our cell typing method consists of three steps (**Supplementary Figure S1)**. First, we select a set of marker genes sufficient for identifying cell types, using previous scRNAseq data. Second, we apply *in situ* sequencing to detect expression of these genes at cellular resolution in tissue sections. Third, gene reads are assigned to cells, and cells to types using a probabilistic model derived from scRNA-seq clusters.

To select a gene panel, we developed an algorithm that searches for a subset of genes that can together identify scRNA-seq cells to their original clusters, after downsampling expression levels to match the lower efficiency of *in situ* data (see Methods). The gene panel was selected using a database of inhibitory neurons from hippocampus^5^ (**Supplementary Figure S2**) as well as isocortex^2^, and the results were manually curated prior to final gene selection. The algorithm identified familiar interneuron markers such as *Sst, Vip, Npy*, and *Cck*, but also many genes identified only by transcriptomic analysis (e.g. *Cxcl14, Ntng1, Id2*). The final gene sets were chosen by manual curation of these lists, excluding genes that were likely to be strongly expressed in all cell types even if at different levels, and favoring genes which have been used in classical immunohistochemistry (**Supplementary Table S1, Supplementary Figure S3**). A further set of three genes were excluded after initial experiments, as their expression was widespread in neuropil and did not help identify cell types (*Slc1a2, Vim, Map2*). The final panel contained 99 genes.

To generate RNA expression profiles, we modified the barcode-targeted *in situ* sequencing method described by Ke *et al*^13^ (**Supplementary Figure S4**). A library of padlock probes was generated to match the selected genes. For each gene, we selected a set of target sequences along the exon sequence, such that the two arms of each padlock probe together match a 40-basepair sequence along on the mRNA. In addition to the recognition arms, each padlock probe contained a 4-basepair barcode, an “anchor sequence” allowing all amplicons to be labelled simultaneously, and a 20-basepair hybridization sequence allowing for additional readouts. For more weakly expressed genes, we designed probes matching multiple target sequences along the mRNA length, which aided their detection without compromising detection of others (**Supplementary Figure S5**). In total we designed 755 probes for 99 genes, but used only 161 barcodes out of 1024 (= 4^5^) possible combinations to allow error correction (for probe sequence and barcodes see **Supplementary Table S2)**.

To apply the method to a tissue section, mRNA is enzymatically converted to cDNA *in situ* and then degraded. The padlock probe library is applied, and a ligation enzyme selectively circularizes only probes perfectly matching their cDNA target sequences. The circularized probes are rolling-circle amplified (RCA), generating sub-micron sized DNA molecules (rolling-circle products: RCPs), each carrying hundreds of copies of the probe’s barcode. The RCP barcodes are identified with a standard fluorescence microscope (20x objective) in five rounds of multi-color imaging (**Figure 1A**). Finally, RCPs for two genes which express so strongly their signal would swamp others *(Sst* and *Npy*) are detected separately in a 6^th^ round by hybridizing fluorescent probes to their target recognition sequences. Data is analyzed using a custom pipeline, including point-cloud registration to deal with chromatic aberration in the images, and compensation for optical or chemical crosstalk between bases in the sequencing readout (**Figure 1B; Supplementary Figure S6** and Methods). These improved chemical and analytic methods achieved a density of reads sufficient for fine cell type assignment.

**Figure 1.**
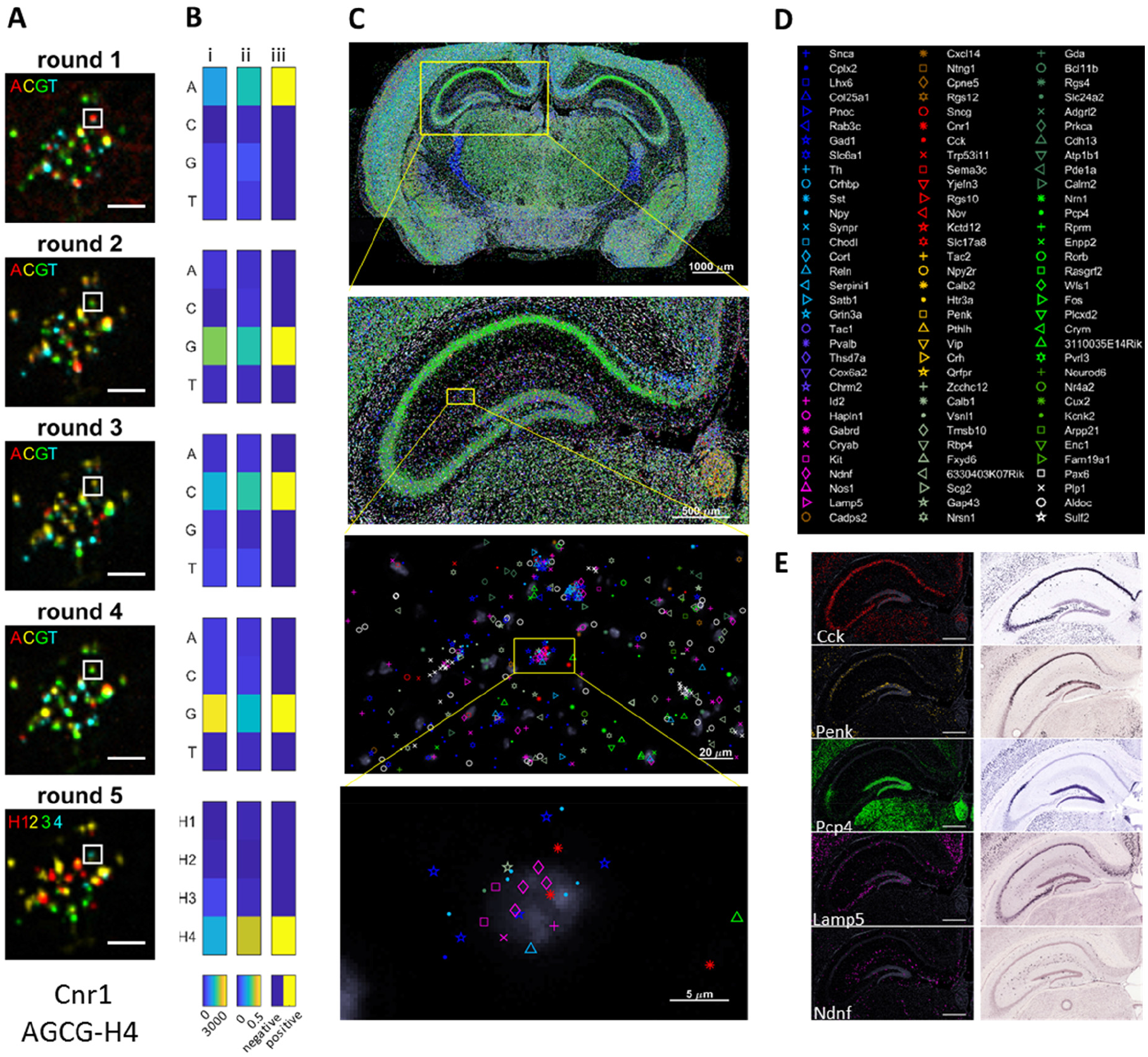
Molecular maps. **A)** Pseudocolor images showing barcode sequencing readout for a region corresponding to one cell. Top to bottom, base-specific fluorophores in the four cycles of sequencing by ligation, and for the fifth cycle of barcode specific hybridization. The white square shows a single RCP of barcode AGCG-H4. Scale bars: 5 μm. **B)** Gene-calling for this RCP. Left: pseudocolor representation of raw fluorescence intensities; Middle, intensity after crosstalk compensation; Right, best fit barcode (AGCG-H4, encoding the gene *Cnr1*). **C)** Distribution of 99 genes at different zoom levels. From top to bottom: a complete coronal mouse brain section; left hippocampus; the border of stratum radiatum and stratum lacunosum moleculare; finally, zoom-in to reads for the cell whose raw fluorescence is shown in panel (A). **D)** Code symbols for the 99 marker genes. **E)** Comparison of the distribution of five markers in the hippocampus as determined by pciSeq (left column) with the distribution shown in the Allen Mouse Brain Atlas (right column). Scale bars: 500 μm.

Our first experiments were performed using probes against a subset of 84 genes on four coronal sections of mouse brain (10 μm fresh frozen). After verifying that detected expression patterns match *in situ* hybridization data from the Allen Mouse Brain Atlas^21^, we continued with two further experiments using the full 99-gene panel, on two and eight coronal sections, respectively. All fourteen sections were from one P25 male CD1 mouse and covered different parts of the dorsal hippocampus (**Supplementary Figure S7**). Each section contained roughly 120,000 cells and in total 15,424,317 reads passed quality control (**Supplementary Table S3)**. To display reads corresponding to 99 genes on a single map, we displayed each read with symbols whose colors grouped genes often expressed by similar cell types, and glyph distinguished genes within these color groups (**Figure 1, C and D**).

Expression patterns were consistent with expectation at multiple levels of detail. At the whole-brain level (**Figure 1C**, top), differences between regions could be seen, for example with the inhibitory thalamic reticular nucleus dominated by inhibitory-associated genes (blue) and the CA1 pyramidal layer dominated by pyramidal-associated genes (green). Zooming in to the hippocampus (**Figure 1C**, 2^nd^ row) revealed differences between cell layers, for example with stronger expression of genes associated with *Sst* neurons (cyan) in stratum oriens, neurogliaform-associated genes (pink) in stratum lacunosum-moleculare, and non-neuronal genes (white) in the white matter. Zooming further to single neurons (bottom two rows) showed genes that appeared to be grouped together in combinations as expected from scRNA-seq. Expression patterns of all genes present in the Allen Mouse Brain Atlas^21^ matched at a corresponding coronal level (examples in **Figure 1E**). Read densities were consistent between experiments, even with different gene panels, further supporting the reliability of the technique (r = 0.93; **Supplementary Figure S8A**).

We focused our cell typing analysis on hippocampal area CA1 (**Supplementary Figure S9**), where we can use the known laminar locations of several identified cell types to validate the method. We used the approach to identify the types of 27,338 cells typed as CA1 neurons, from 28 hippocampi. Data files for all experiments are available at https://figshare.com/s/88a0fc8157aca0c6f0e8, and an online viewer showing reads and probabilistic cell type assignments is at http://insitu.cortexlab.net.

A fundamental challenge for *in situ* cell typing is assigning genes to cells. mRNA molecules can be found throughout a cell’s cytoplasm, and boundaries between cells are difficult to obtain in 2D imaging. In the current approach, we counterstained all sections with DAPI to reveal cellular nuclei. Applying standard watershed segmentation to these cells yielded boundaries that contained many, but not all the genes that manual inspection suggested belonged to them (**Figure 2A**). To solve this problem, we developed an algorithm which leverages scRNA-seq data to assign genes to cells more accurately than would be possible by spatial location alone (**Figure 2A**, straight lines). The model uses scRNA-seq clusters to define an expected read density for each gene in each cell type, and assigns a prior probability of assigning reads to cells based the DAPI image. It employs a variational Bayesian algorithm to compute a posterior probability distribution for the cell assigned to each read, and the type assigned to each cell. Note that the algorithm does not take into account a cell’s laminar location, allowing this to be used for validation.

**Figure 2.**
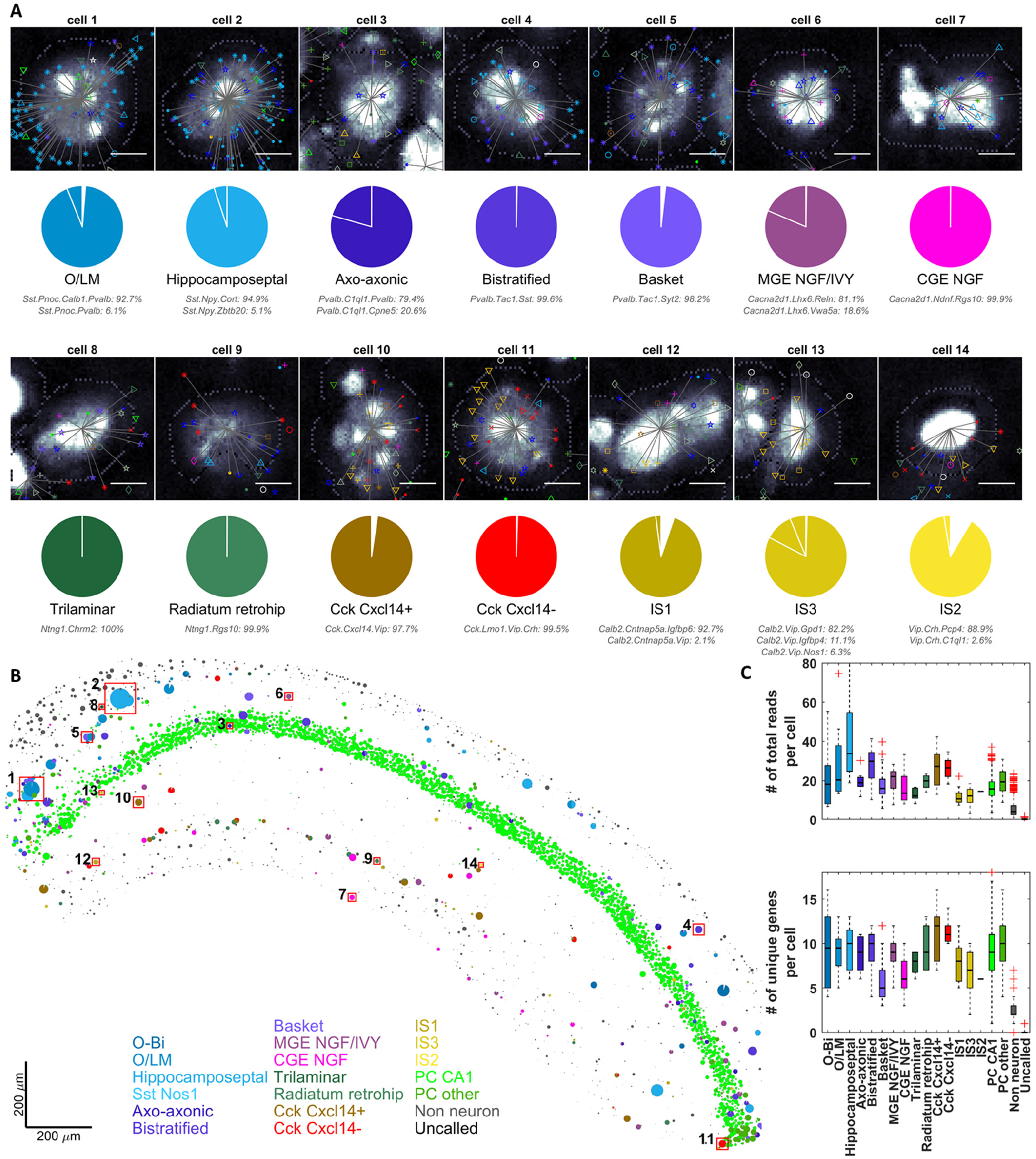
Cell type map of CA1 from an example experiment (experiment 4–3 right hemisphere). **A)** Reads are assigned to cells, and cells to classes using a probability model based on scRNA-seq data. Top row: distribution and assignment of reads for fourteen example cells. Colored symbols indicate reads (color code as in **Figure 1D**). Grayscale background image indicates DAPI stain with watershed segmentation as dotted line. Straight lines join reads to the cell for which are assigned highest probability. Scale bars: 5 μm. Bottom row: pie charts showing probability distribution of each class for the same example cells. Colors indicate broad cell types; segments show probabilities for individual scRNA-seq clusters (named underneath). **B)** Spatial map of cell types across CA1. Cells are represented by pie charts, with size proportional to the number of reads assigned to the cell. Numbers identify the example cells in A. **C)** Box-and-whisker representation of total read count per cell of each type (top) and average number of unique genes per cell of each type (bottom).

To represent the results on a spatial map, we displayed each cell’s class assignments by a pie-chart, of size proportional to total gene count, and with the angle of each color-coded slice indicating the probability of assignment to that class (**Figure 2B;** see also **Supplementary Figure S10;** for all cell type maps, see **Supplementary Appendix;** online viewer at http://insitu.cortexlab.net). The algorithm was able to confidently identify 16 inhibitory classes: 3 types of interneuron-selective cell; 2 types of Cck cell; 2 types of neurogliaform cell; 2 types of GABAergic projection cell; 3 types of parvalbumin cell and 4 types of somatostatin cell (**Supplementary Tables S4 and S5**). Further subdivisions of these types, identified by scRNA-seq clusters, were called with varying levels of confidence. Although our panel was primarily aimed at distinguishing inhibitory neurons, we also obtained confident distinction of two types of pyramidal cell, as well as nonneuronal cells. The probabilistic algorithm allows diagnostics which show which genes provided evidence for calling as one type over another (**Supplementary Figure S11**). The average number of gene reads per cell was over 20 for most targeted cell types, and the number of unique genes detected per cell was in the range 5 to 10 (**Figure 2C**).

The algorithm’s cell type assignments conformed closely to known combinatorial patterns of gene expression in fine CA1 inhibitory subtypes. For example, the identification of *Sst* cells as O/LM or hippocamposeptal correlated with further expression of *Reln* or *Npy*^22, 23^ (**Figure 2A**; cells 1,2); identification of *Pvalb* cells as axoaxonic, basket or bistratified correlated with further expression of *Pthlh, Satb1/Tac1*, or *Sst/Npy*^6, 22, 24^ (Cells 3–5); identification of neurogliaform cells as CGE-derived or MGE-derived/Ivy correlated with further expression of *Ndnf/Kit/Cxcl14* or *Lhx6/Nos1*^2, 5, 25,26^; identification of long-range projection neurons as trilaminar or radiatum-retrohippocampal correlated with expression of *Chrm2* or *Ndnf*/*Reln*^23, 27^(Cells 8,9)*; Cck* cells were identified as two subtypes correlated with expression of *Cxcl14*, with both subtypes expressing *Cnr1* and further subdivided by *Vip* expression ^5, 28, 29^(Cells 10–11); finally, inhibitory-selective cells were divided into three classes correlated with the combinatorial expression of *Calb2* and *Vip* ^30, 31^ (Cells 12–14). Across all experiments, the patterns of both classical and novel interneuron markers were consistent with scRNAseq results, as well as the known biology of CA1 interneurons (**Supplementary Figure S12**). Moreover, the cell type composition was consistent between the left and right hemispheres **(Supplementary Figure S8B**).

As the pciSeq algorithm did not use cell’s laminar location to determine its class assignment, we were able to validate both the *in situ* cell typing method, and the scRNA-seq classification it relies on, by verifying that cell classes it identifies are found in appropriate layers. The layers in which cell types were identified were consistent with known ground truth (**Figure 3**). Cells identified as O/LM, hippocamposeptal, or O-Bi showed a strong preference for their known locations in *stratum oriens* (*so*), while neurons identified as bistratified also could be found in *stratum pyramidale* (*sp*)^32,33^ (Sst/Nos1 cells were too rare to reliably localized; **Supplementary Figure S13**). Cells identified as basket cells were found in *sp* and less often *so*, while rarer cells identified as axo-axonic were found in the pyramidal layer nearly exclusively^34^. Amongst cells identified with neurogliaform cells, those identified as having developmental origin in the medial ganglionic eminence (including Ivy cells) were found throughout all layers, while those identified as having origins in caudal ganglionic eminence were found nearly exclusively in *stratum lacunosum-moleculare* (*slm*), consistent with previous reports^25,26^. Of the two transcriptomic classes identified with long-range projecting GABAergic neurons, those identified as trilaminar cells were located primarily in *so*^23, 35, 36^; while those identified as radiatum retrohippocampal were found at the border of *stratum radiatum*(*sr)* and *slm*^23,27, 37,38^. Cck interneurons were divided into two primary classes, with the *Cxcl14*-positive class located primarily in *sr*, close to the *slm* border, and the *Cxcl14-*negative class in all layers, as previously predicted^5^. Amongst interneuron-selective subtypes, cells identified as IS1 were found in all layers as expected^31^, while IS3 cells were located primarily in *sp* and *sr*, but very rare in *slm*^28^ (IS2 cells were too rare for reliable quantification of their laminar distribution). Reassuringly, cells identified as CA1 pyramidal were found exclusively in *sp*, while cells identified as non-CA1 excitatory could also be occasionally found in *so*, with the latter reflecting primarily subicular pyramidal cells, which can also be found extending into *so* at the CA1/subiculum boundary. Thus, our method identified 18 types of CA1 neuron with the expected laminar distributions, both confirming the predictions of our cell type identifications, and the *in situ* sequencing method itself.

**Figure 3.**
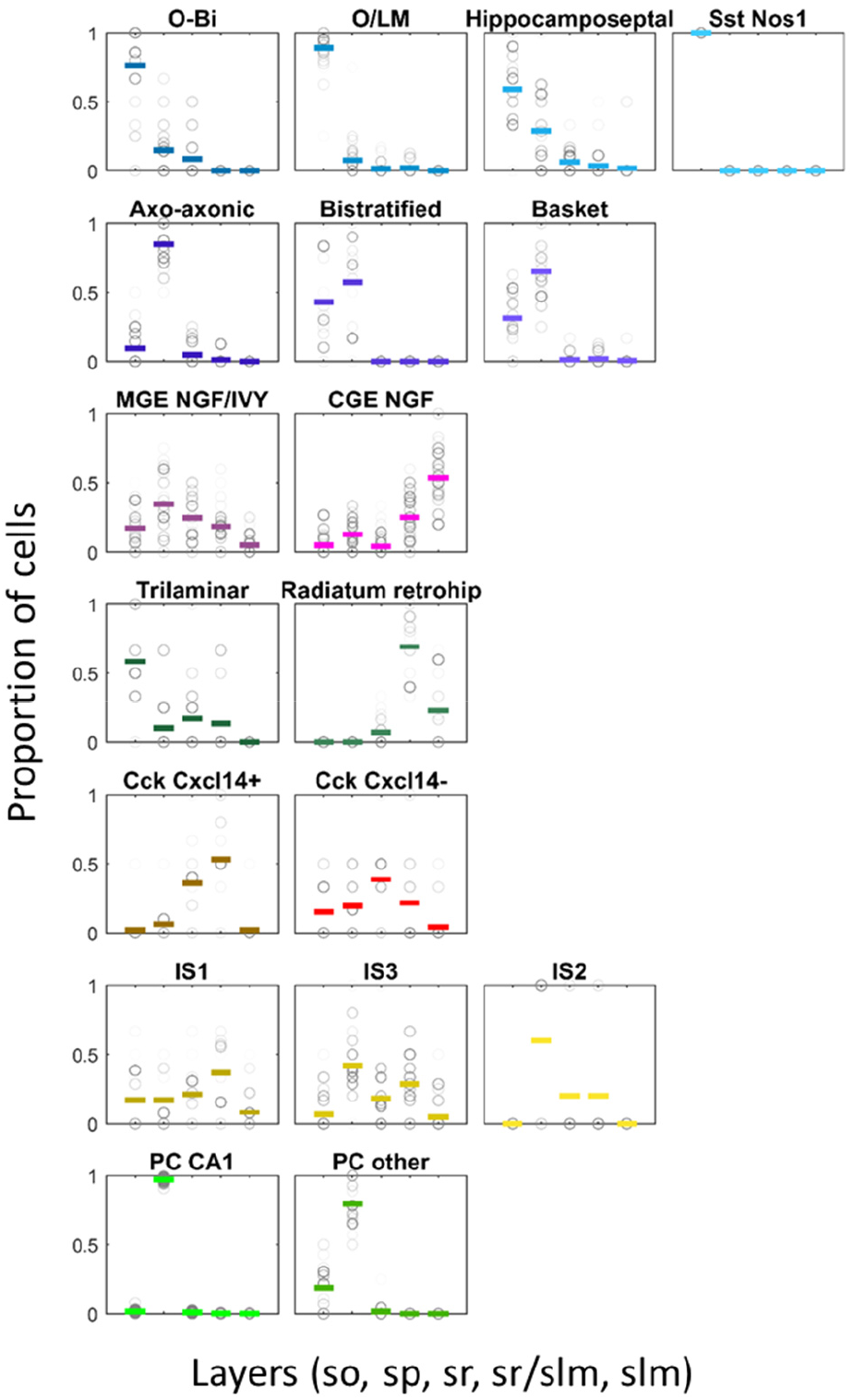
Fraction of each cell class found each CA1 layer. Circles indicate means of a single experiment with gray level representing number of cells of that class in the experiment; colored lines denote grand mean over all 28 hippocampi. In each plot, the 5 x-axis positions represent layers: stratum oriens (so), stratum pyramidale (sp), stratum radiatum (sr), border or strata radiatum and lacunosum-moleculare (sr/slm); stratum lacunosummoleculare (slm).

To conclude, we have shown it is possible to identify closely related cell classes *in situ* by levering information from scRNA-seq to design a probe panel and cell identification algorithm for *in situ* sequencing. A key to this method’s success is its very low false-positive gene detection rate. Indeed, while the 16 inhibitory classes identified by this algorithm closely match known biology, a recent SNAIL-based method^15^ with higher total read density but also possibly higher erroneous reads identified just four or five classes, which were dominated by single genes and do not match combinatorial expression patterns suggested by scRNA-seq and classical analyses. We found that sixteen classes of CA1 inhibitory cell could be identified with just 99 genes chosen using scRNA-seq data. Large-scale scRNA-seq projects are now underway for the whole body, and the data required to design panels and apply this method to all tissues will soon be available. The pciSeq approach requires only low-magnification imaging (in contrast to FISH-based methods), and so may be applied high throughput, raising the possibility of body-wide spatial cell type maps in the near future.

## Acknowledgements

This work was supported by grants from the Wellcome Trust (108726, to KDH, JHL, and MN) and Chan-Zuckerberg Initiative (182811 to KDH). We thank Peter Somogyi, Matteo Carandini, Sten Linnarsson, Nicoletta Kessaris and Lorenza Magno for valuable discussions. We thank Kasper Karlsson for visualizing scRNA-seq reads and padlock probe targets in genome browser.

## Author contributions

XQ wrote DNA probe design software, performed experiments, analyzed data, designed in situ sequencing protocol, prepared figures, wrote manuscript. KDH designed and wrote analysis software, wrote manuscript. TH designed in situ sequencing protocol. DN designed and wrote online web viewer. AMM prepared tissue samples. NS contributed to gene panel selection. JHL supervised tissue sample collection. MN designed in situ sequencing protocol, supervised experiments, wrote manuscript.

